# Analysis of Differential Gene Expression and Core Canonical Pathways involved in the Epithelial to Mesenchymal Transition of Triple Negative Breast Cancer Cells by Ingenuity Pathway Analysis

**DOI:** 10.1101/2023.04.07.536005

**Authors:** Santanu Bhattacharya, Somiranjan Ghosh, Hirendra Banerjee

## Abstract

Triple Negative Breast Cancer (TNBC) is a malignant form of cancer with very high mortality and morbidity. Epithelial to Mesenchymal Transition (EMT) is the most common pathophysiological change observed in cancer cells of epithelial origin that promotes metastasis, drug resistance and cancer stem cell formation. Since the information regarding differential gene expression in TNBC cells and cell signaling events leading to EMT is limited, this investigation was done by comparing transcriptomic data generated by RNA isolation and sequencing of a EMT model TNBC cell line in comparison to regular TNBC cells. RNA sequencing and Ingenuity Pathway Software Analysis (IPA) of the transcriptomic data revealed several upregulated and downregulated gene expressions along with novel core canonical pathways including Sirtuin signaling, Oxidative Phosphorylation and Mitochondrial dysfunction events involved in EMT changes of the TNBC cells.

## NTRODUCTION

Triple-negative breast cancer (TNBC) accounts for about 10-15% of all breast cancers[1]. The term triple-negative breast cancer refers to the fact that the cancer cells don’t have estrogen or progesterone receptors (ER or PR) and don’t make any or too much of the protein called human epidermal growth factor receptor-2 (HER2) [1]. These cancers tend to be more common in women younger than age 40, who are Black, or who have a BRCA1 mutation [2]. TNBC differs from other types of invasive breast cancer in that it tends to grow and spread faster, has fewer treatment options, and tends to have a worse prognosis [3, 4]. Triple-negative breast cancer (TNBC) is considered an aggressive cancer because it grows quickly, is more likely to have spread at the time it’s found and is more likely to come back after treatment than other types of breast cancer[1, 3, 4]. Triple-negative breast cancer has fewer treatment options than other types of invasive breast cancer. This is because the cancer cells do not have the estrogen or progesterone receptors or enough of the HER2 protein to make hormone therapy or targeted HER2 drugs work[1]. Hormone therapy and anti-HER2 drugs are not choices for women with triple-negative breast cancer, there is limited scope of chemotherapy; the overall survival rate of TNBC currently is around 77% (American Cancer Society) [4].

Although epithelial-to-mesenchymal transition (EMT) and mesenchymal-to-epithelial transition (MET) have been implicated in the incidence of cancer metastasis and drug resistance, their impact in cancer progression and patient survival is not fully understood. During EMT, epithelial cells lose their polarity, as well as their cell-cell adhesions, and acquire the motile and invasive characteristics of mesenchymal cells [5]. Proteins such as vimentin (VIM) intermediate filament (IF) are generally upregulated when the cell is in the mesenchymal relative to the epithelial status [6].

The VIM RFP reporter cell line (ATCC HTB-26MET) was created using CRISPR/Cas9 gene editing in the parental MDA-MB-231 breast adenocarcinoma cell line (ATCC HTB-26). HTB-26MET harbors a C-terminal red fluorescent protein (RFP) tag on the vimentin gene. This enables the tracking of the EMT status of cells in vitro by monitoring RFP expression. The integrity of the VIM RFP knock-in has been verified at the genomic, mRNA, and protein level for sequence and expression by scientists at American Type Culture Collection (ATCC, USA).

Since EMT has been implicated for breast cancer metastasis, angiogenesis, drug resistance and eventually cancer stem cell formation [7-9], we in this study investigated the differential gene expression and core canonical pathways involved in the EMT changes in a TNBC -EMT model Vimentin-RFP knock in cell line(ATCC HTB-26MET) in comparison to the same non EMT TNBC MDA-MB-231 breast adenocarcinoma cell line (ATCC HTB-26) by analyzing the transcriptomic data obtained by NGS RNA Sequencing and using the **Ingenuity Pathway Analysis** (IPA) Software as licensed by Qiagen Corporation, USA.

## Materials and Methods

### Cell Culture

MDA-MB-231-VIM-RFP (ATCC CRM-HTB-26) were obtained from the American Type Culture Collection (ATCC, Manassas, Virginia) and cultured in Eagle’s Minimum Essential Medium with the inclusion of 0.01mg/mL of insulin and 10μg/mL blasticidin,10% FBS and antibiotics maintained in a 5% carbon dioxide incubator at 37°C. The MDA-MB-231 also purchased from ATCC was cultured in L-15 medium supplemented with 10% FBS and antibiotics and kept in an incubator at 37°C.

### Fluorescence Imaging

The breast cancer cell lines were split by Trypsinization and grown in 6-well cell culture plates, when confluent, the cells were photographed using an Olympus Fluorescence Microscope using the red filter.

### RNA Isolation and Sequencing

All RNA isolation procedures were conducted according to the manufacturer’s protocol (Signosis LLC, Single Cell RT-PCR Assay Kit, Santa Clara, CA). Between 1,000 and 10,000 cells, as confirmed using a cell counter (Denovix CellDrop Brightfield cell counter, Wilmington, DE), were isolated from cell culture and washed with 200 μL of ice cold 1X PBS. Ice-cold cell lysis buffer (50 μL) was added, and the solution was then snap-frozen at -80°C for 5 min. Cells were incubated on ice for 10 min and centrifuged at 10,000 g for 2 min. Supernatant was transferred to a fresh nuclease-free microcentrifuge tube. DNAse I (1 μL) was added, and the sample was incubated at 37°C in a water bath for 30 min and then inactivated at 75°C for 10 min. Supernatant was then placed in ice and stored at -80°C. The isolated RNA was sent to PrimBio Research Institute LLC (Exton, PA) for RNA transcriptome sequencing. Results obtained from RNA sequincing analysis yielded a fold change which was used to develop pathways using Ingenuity Pathway Analysis (IPA) Software licensed from Qiagen Corporation, USA.

### Ingenuity Pathway Analysis Methodologies

IPA was used to organize and analyze data obtained from RNA sequences. RNA sequences were generated from Prim Bio Research Institute (Exton, PA) with cells cultured in this laboratory. Weight was assigned to certain gene products based on significance and fold change. Relevant pathways and import molecules emerged in the context of complex interrelated cellular processes.

## RESULTS

A fluorescent image was taken of the MDA MB 231 breast cancer cells and Vimentin-RFP tagged knock in EMT MDA-MB-231 cells detecting the constitutively expressed Vimentin gene by red fluorescence. This image (Figure 1) proves that the RFP tagged Vimentin gene which is upregulated in the EMT changes of breast cancer cells is correctly in frame inserted by the CRISPR-CAS9 technique in the genome of these TNBC cells and inducing EMT.

**Figure 1:**
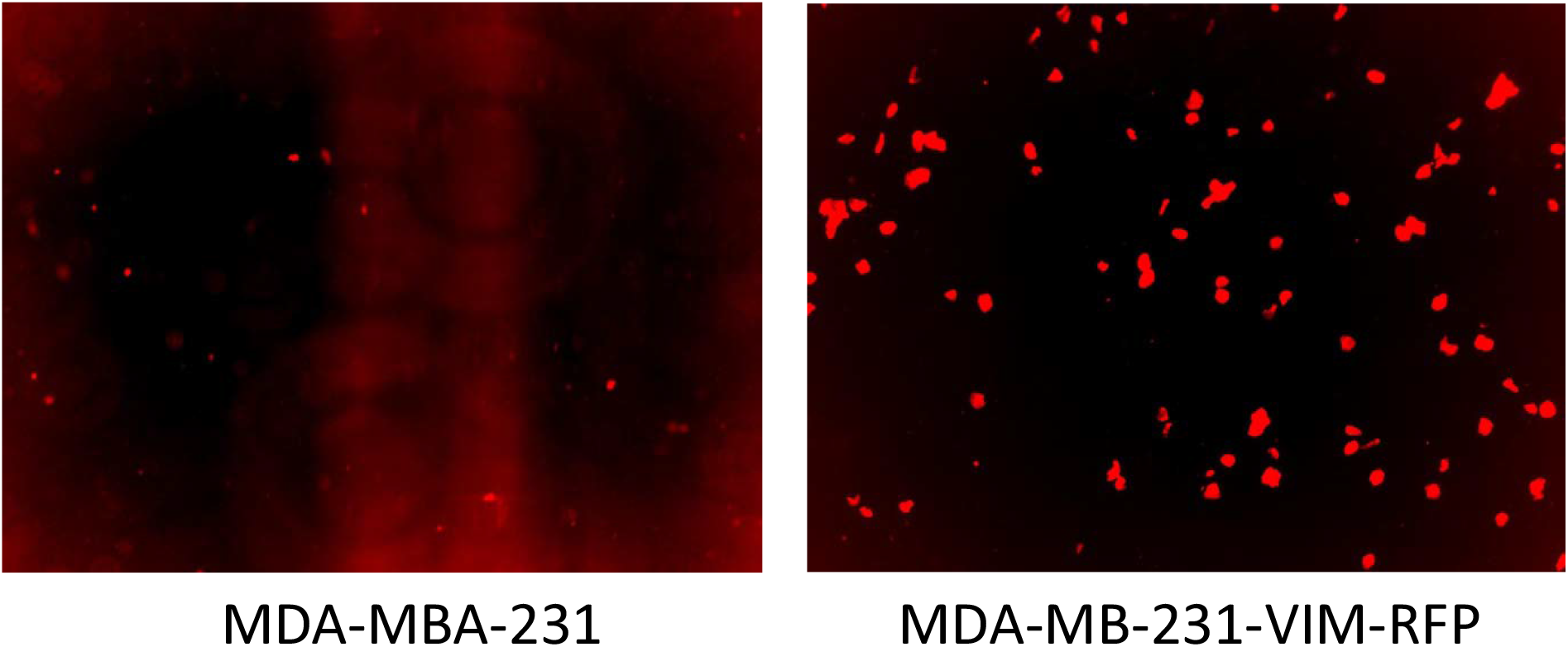
Fluorescent imaging of a non-transgenic breast cancer cell line (left) compared to the genetically engineered breast cancer cell (right). Red fluorescent demonstrates RFP labeled vimentin expression.

The Ingenuity Pathway Analysis software created by Qiagen was used to process the raw complete transcriptome data of the differential gene expression of the MDA-MB-231 -VIM-RFP in comparison to MDA-MB-231 obtained from Primbio. A forecast model of potential upregulated and downregulated key canonical pathways that are important in cell life, cell morphology, and cellular functions of the cells was created by IPA based on the differential gene expression. Since there were several down and up regulated genes identified, we considered the top canonical pathways and taking that data ran a core analysis. The results deciphered three canonical pathways which were the most common overlapping pathway among all the different pathways detected analyzing the transcriptomic data. Figure 2 is Based on a positive z-score, the orange hue indicates that the pathway is upregulated, and a negative z-score with blue color indicates that the pathway is downregulated. The gray hue indicates that the pathway is not activated, and the white color indicates that the activity is uncertain. The three core canonical pathways most frequently observed during the investigation are the Sirtuin Signaling, Oxidative Phosphorylation, and the Mitochondrial Dysfunction pathways. The Sirtuin Signaling pathway had the highest z-score, which helped to analyze the rest of the pathways.

**Figure 2:**
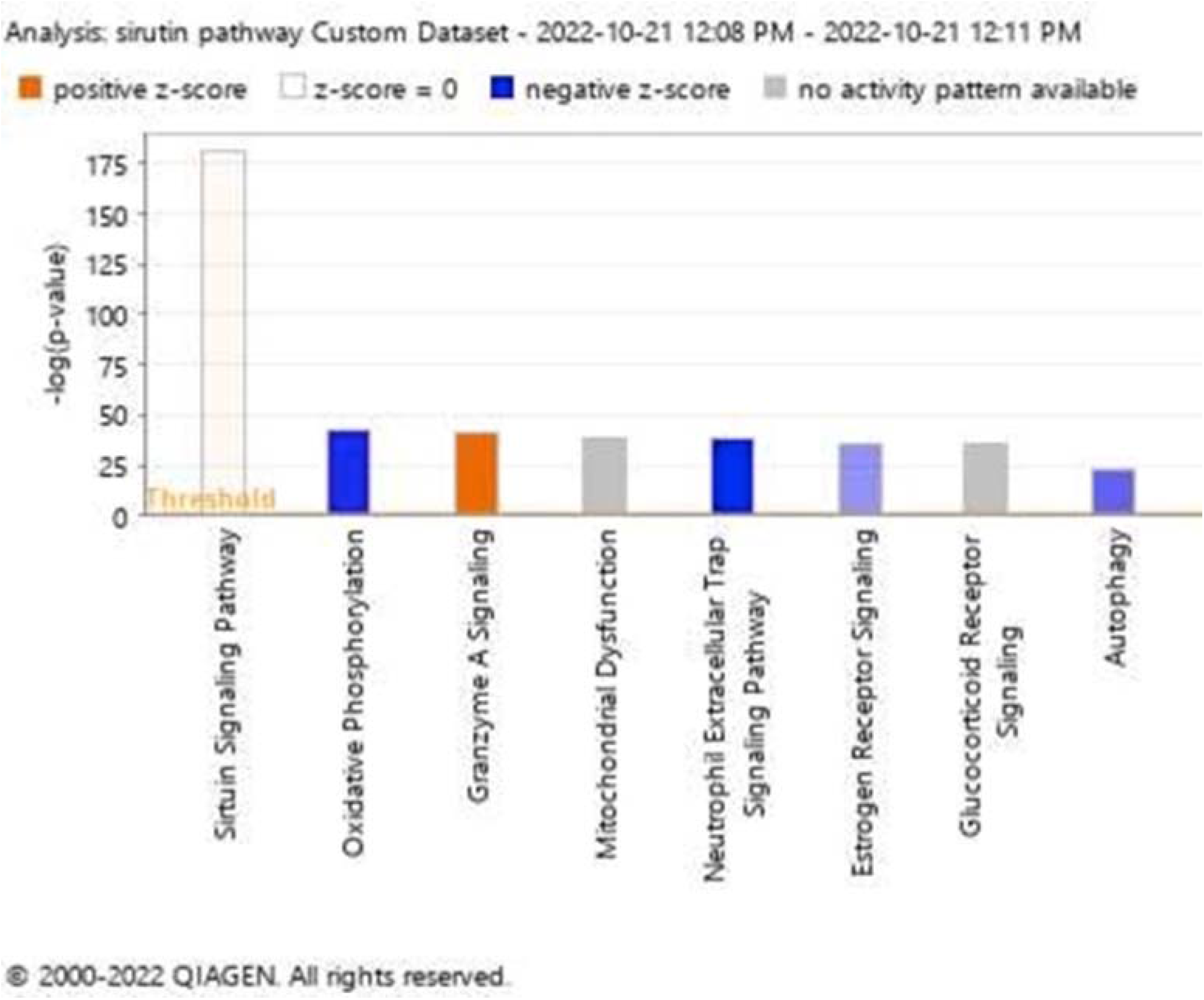
Based on IPA software analysis of RNA sequencing data. This figure shows the primary canonical pathways involved in EMT MDA-MB-231 vs regular breast cancer cell line. This graph displays two pathways that may be up- and down-regulated.

These fundamental canonical pathways play a part in several processes and the development of diseases. There are a few routes that involve cancer genesis, proliferation, and cell survival in the setting of cancer progression, Figure 3 is a bubble chart indicating the predicted canonical pathway due to gene regulation in the comparison of EMT model MDA-MB-231 vs regular MDA breast cancer cells. The sizes of the bubbles states how many genes are associated with that specific pathway. The Sirtuin Signaling is involved in cellular immune response, cellular stress, injury, and many more cellular events as shown in Figure 4. Table 1 and 2 show the differentially expressed genes as identified from the transcriptomic data, whereas Table 3 and 4 describe the functionality of those differentially expressed genes in various disease process including cancer.

**Table 1:**
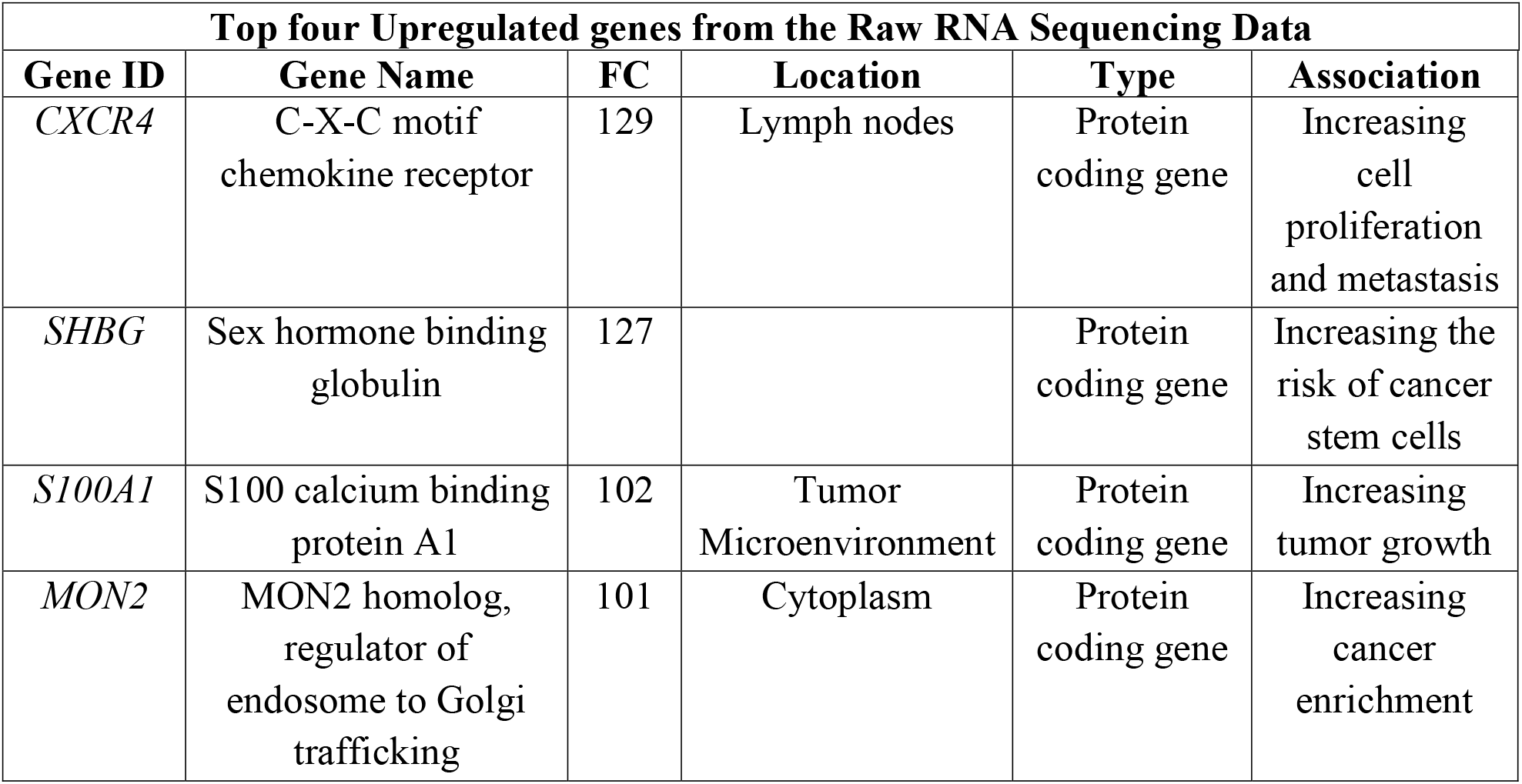
Raw transcriptomic RNA sequencing data. The top four genes with the largest fold change (FC) upregulation in the EMT MDA-MB-231 vs. ordinary MDA breast cancer cell line are represented in this data. In breast cancer, CXCR4 exhibits the highest elevated fold change. Three out of the four identified as an oncogene which are CXCR4, SHBG, S100A1.

**Table 2:** RNA sequencing full transcriptome raw data. This information shows which four genes were downregulated by the greatest amount in EMT MDA-MB-231 breast cancer cells as compared to normal MDA breast cancer cells.

| **Top four Downregulated genes from the Raw RNA Sequencing Data** |  |  |  |  |  |
| --- | --- | --- | --- | --- | --- |
| **Gene ID** | **Gene Name** | **FC** | **Location** | **Type** | **Association** |
| *GNB2L1* | Guanin nucleotide-binding protein subunit beta-2-like 1 | -12.7 |  | Protein Coding | Downregulate metastasis |
| *ECHS1* | Enoyl-CoA hydratase, short chain 1 | -8.5 |  | Protein coding | Enhances PP2 induced apoptosis in breast cancer |
| *YBX1* | Y-box binding protein 1 | -8.5 |  | Protein coding | Decrease the response to tamoxifen and fulvestrant |
| *CSTB* | Cystatin B | -8.4 |  | Protein Coding | Regulates malignant |

**Table 3:** Disease and Function analysis of upregulated genes associated with breast cancer proliferation in MDA-MB-232 vs regular MDA breast cancer cell line. In the comparison of Sirtuin Signaling, Oxidative Phosphorylation, and Mitochondrial Dysfunction pathway.

| **Downregulated Genes Associated with Breast Cancer Cell Proliferation in the comparison of genes in Sirtuin Signaling, Oxidative Phosphorylation, and Mitochondrial Dysfunction pathway** |  |  |  |  |  |  |  |
| --- | --- | --- | --- | --- | --- | --- | --- |
| **Gene ID** | **Gene Name** | **Log Ratio** | **Location** | **Family** | **Association** | **Known Function** | **Data set** |
| *ATP5PF* | *ATP synthase peripheral stalk subunit F6* | *-4.7* | *Cytoplasm* | *Transporter* | *Cell migration* | *Increase proliferation* | *Down regulated* |
| *MAPK12* | *Mitogen-activated protein Kinase 12* | *-5.1* | *Cytoplasm* | *Kinase* | *Cancer stem cells* | *Increase proliferation* | *Down regulated* |

**Table 4:**
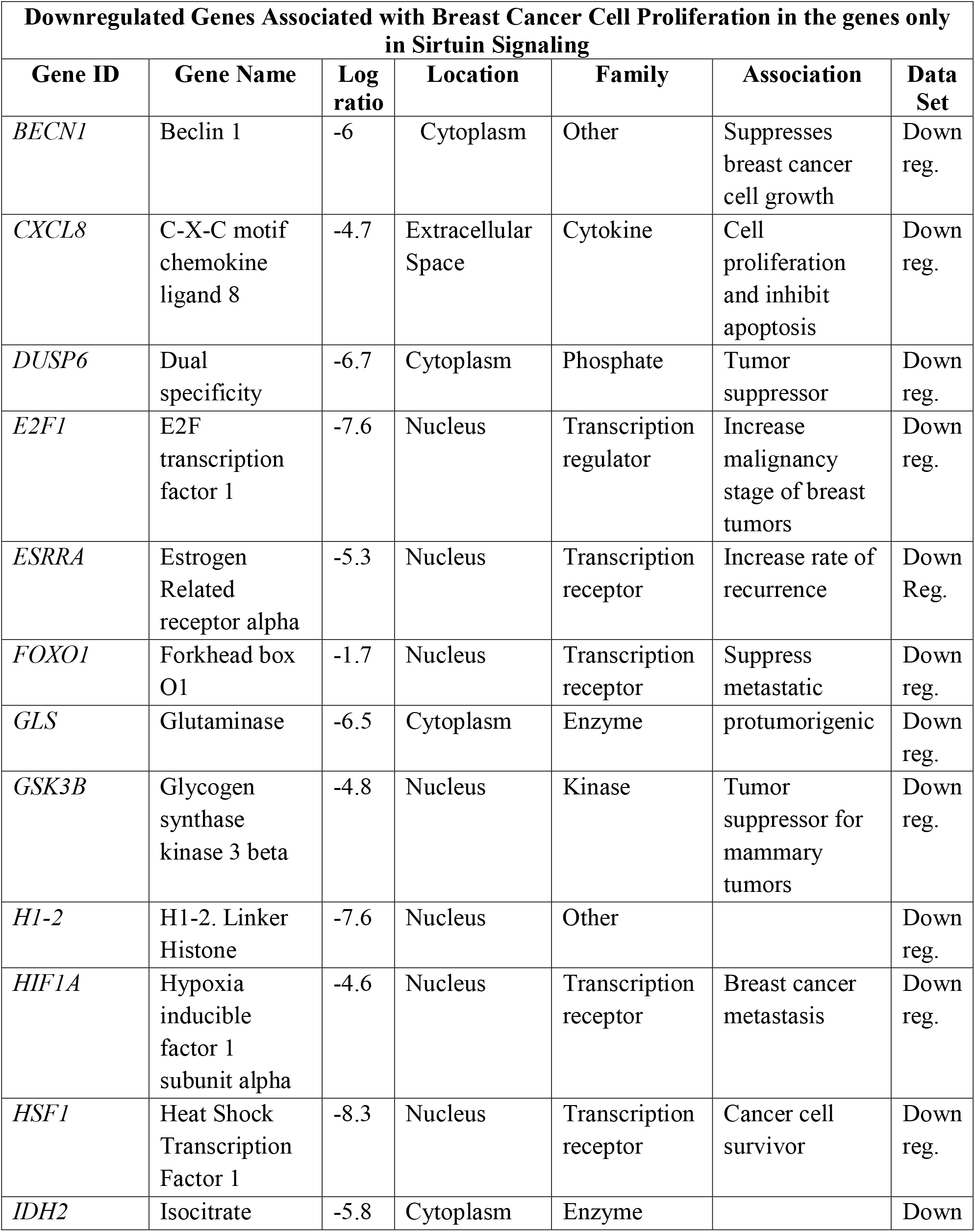
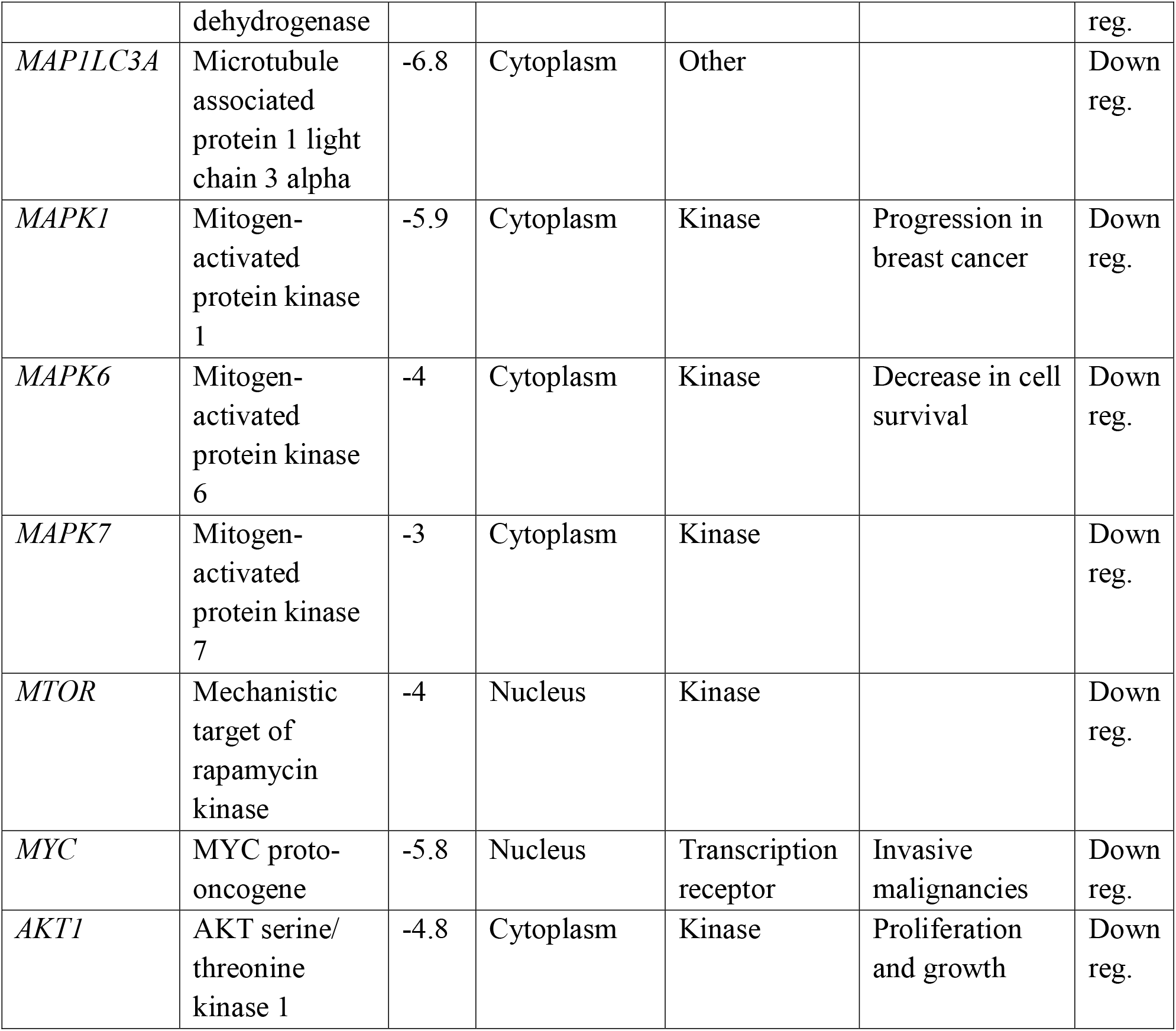
Disease and Function analysis of downregulated genes associated with breast cancer proliferation in MDA-MB-232 vs regular MDA breast cancer cell line associated with the Sirtuin Signaling Pathway.

**Figure 3:**
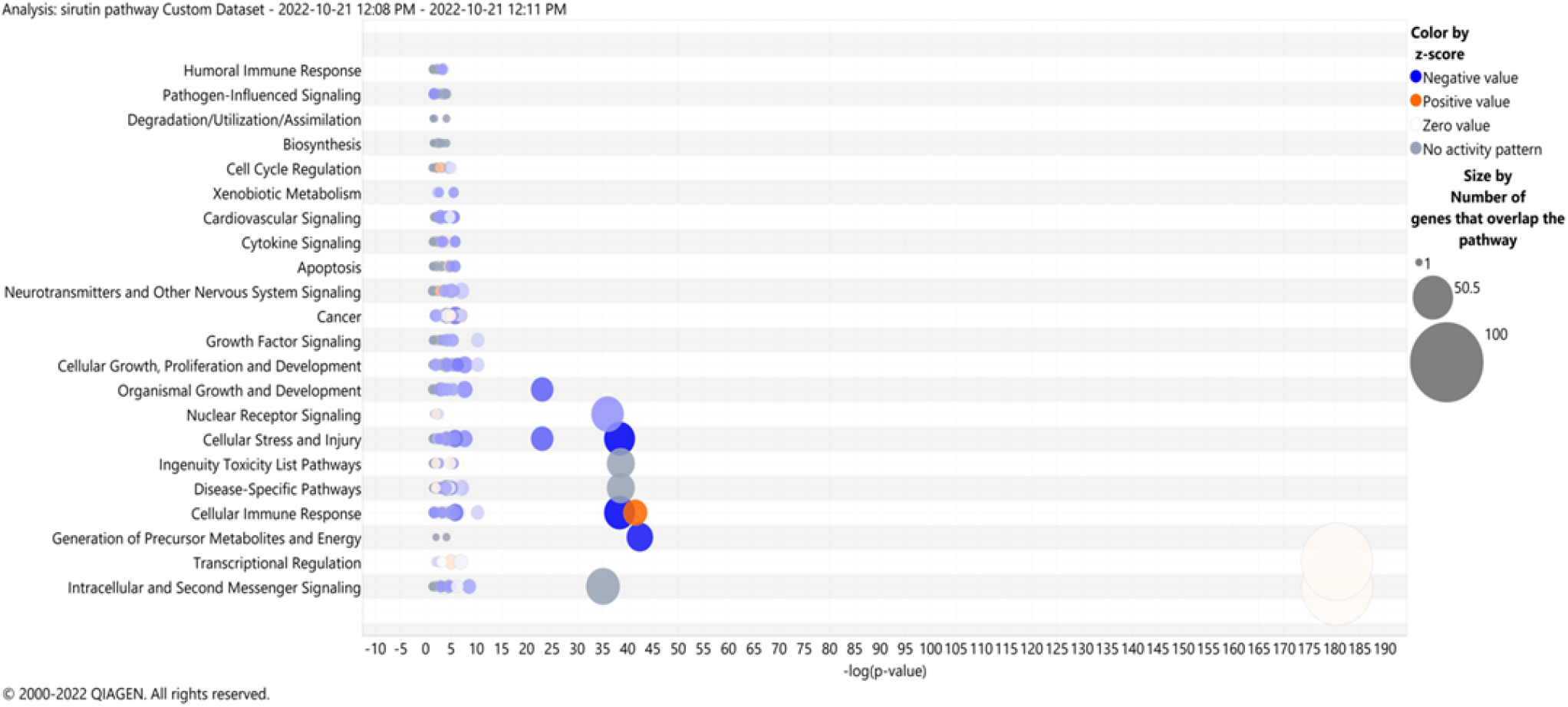
Based on the log fold change activation score of the canonical pathways of MDA-MB-231 vs. MDA-MB-231 EMT breast cancer cell line, this bubble chart displays the expected known canonical pathways gene regulation in regular MDA-MB-231 versus MDA-MB-231 EMT breast cancer cells in the context of function and disease development. The circles’ size reflects the number of genes involved in the pathway, although the color coding is the same as that of the canonical pathway bar graph.

**Figure 4:**
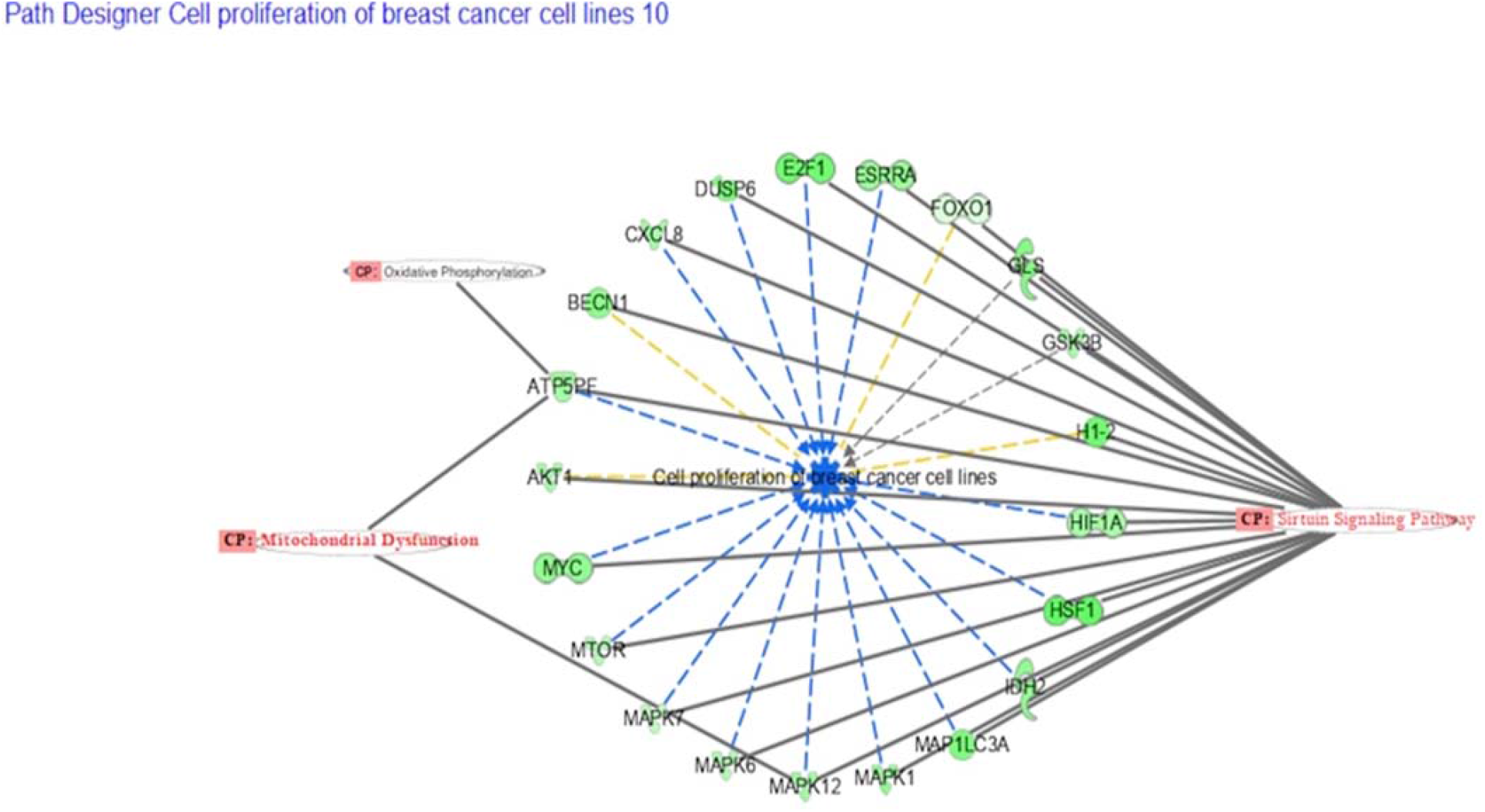
In the IPA data software the disease and function analysis of the RNA sequencing data from Primbio. Each one of these genes are known to be involved in cellular proliferation of breast cancer. These are the top three canonical pathway that are involved with cell proliferation with genes in common. The comparison of each gene and its known function is stated, in this case all the genes are decreasing cell proliferation (Table 3)

## DISCUSSION

The main purpose of this investigation was to study the differential gene expression in the EMT model of MDA-MB-231-VIM-RFP comparison to regular MDA-MB 231 breast cancer cell line. The IPA software played an important role in analyzing results, taking the large transcriptomic data obtained by RNA Sequencing. This detail analysis identified three main core canonical pathways, along with the differentially expressed genes and their role in cancer and other diseases. As evident from literature, these identified genes are highly associated in growth, EMT and metastasis of various cancers including breast cancer. We will outline their significant role in cancer development in the following section of the manuscript.

Regarding the upregulated genes, C-X-C chemokine receptor type 4 (CXCR4) is a G protein-coupled receptor (GPCR) that is expressed on the surface of various cell types, including immune cells, cancer cells, and nervous system cells[10, 11]. It binds to its ligand, CXCL12 (also known as stromal cell-derived factor 1 or SDF-1) and activates intracellular signaling pathways that regulate various cellular processes such as cell proliferation, survival, migration, and differentiation[12]. Some cancer therapies target CXCR4 to inhibit the growth and spread of cancer cells[13]. In cancer, CXCR4 has been found to be overexpressed in a variety of human tumors, including breast, ovarian, and lung cancer[13]. The binding of CXCL12 to CXCR4 on cancer cells promotes their growth, survival, and migration, which contributes to cancer progression and metastasis[10-13]. Therefore’, CXCR4 is considered a potential therapeutic target for cancer treatment. Some drugs that target CXCR4 have been developed for cancer treatment’, such as plerixafor (AMD3100), which prevent the binding of CXCL12 to CXCR4 and thus inhibit the growth and spread of cancer cells [14]. These drugs have shown promising results in preclinical studies and are currently being evaluated in clinical trials for the treatment of various types of cancer [15]. Sex hormone-binding globulin (SHBG) is a protein, that binds to the sex hormones testosterone and estrogen in the bloodstream and is produced in the liver and regulates the levels of these hormones by binding to them and preventing them from interacting with target cells [16, 17]. Elevated levels of SHBG have been observed in certain types of cancer, including prostate cancer, breast cancer, and ovarian cancer [18-20]. Research suggests that SHBG may play a role in the progression of these cancers by regulating the levels of hormones that promote cell growth[18-20].

S100A1 is a multifaceted protein that plays a role in various physiological processes, particularly in muscle physiology and has been proposed as a potential therapeutic target for various pathological conditions related to calcium homeostasis disturbances, muscle and neuronal diseases [21, 22]. S100A1 has been found to be involved in the development and progression of cancer[23]. Studies have shown that S100A1 is overexpressed in various types of cancers, including breast cancer, melanoma, osteosarcoma and prostate cancer [23-26]. The overexpression of S100A1 in cancer cells has been linked to several hallmarks of cancer, such as cell proliferation, survival, invasion, and metastasis [23, 24]. In breast cancer, S100A1 has been found to be overexpressed in invasive ductal carcinomas, which are the most common type of breast cancer [27]. Studies have shown that S100A1 promotes the proliferation and invasion of breast cancer cells and is associated with a poor prognosis in breast cancer patients [27, 28]. MON2 is a protein that belongs to the family of Mon1-Ccz1 complex, which is responsible for recruiting the small GTPase Rab9 to endosomes, which is an essential step in the transport of endosomal cargo to the Golgi [29]. There is limited research on the role of MON2 in cancer, but some studies have suggested that MON2 may play a role in the development and progression of certain types of cancer. One study has shown that MON2 is overexpressed in breast cancer cells and is associated with a poor prognosis in breast cancer patients. The study suggests that MON2 may promote the proliferation and invasion of breast cancer cells by regulating endosome-to-Golgi trafficking [30].

The analysis of the down regulated genes showed guanine nucleotide-binding protein subunit beta-2-like 1 (GNB2L1) protein also known as receptor for activated protein kinase C1 (RACK1), that belongs to the family of G protein, is involved in cell proliferation, migration and chemoresistance [31, 32]. Short chain enoyl coenzyme A hydratase 1 (ECHS1) protein is an enzyme that belongs to the fatty acid metabolic pathway [33]. This gene has been shown to be involved in colon and breast cancers [34-36]. Y-box protein (YBX1) is a transcription factor that binds to specific DNA sequences called Y boxes and regulates gene expression. It has been found to be involved in several cellular processes including growth differentiation and stress response [37]. This gene has been shown to be involved in breast, colon and lung cancer and involved in chemo resistance and formation of cancer stem cells [38, 39]. Cystatin B (CSTB) is a cysteine protease inhibitor that is poorly expressed in lung and colon cancers with poor prognosis [40].

Silent information regulation factor 1 (sirtuin Type 1, SIRT1), as a kind of NAD+ dependent class III histone deacetylation enzyme, has been found to be involved in tumor proliferation, invasion, and metastasis. The roles of SIRT1in breast cancer is multifaceted depending on its substrate from upstream or downstream signaling pathway, overexpression of SIRT1 significantly promoted breast cancer growth both in vitro and in vivo, whereas knockdown of SIRT1 inhibited these phenotypes. Furthermore, SIRT1 was found to interact with Akt directly, consequently promoting the activity of Akt in breast cancer cells in vitro and positively correlating with expression of Akt, P-Akt, in breast cancer tissues in vivo [41].

Mitochondria have been implicated in cell transformation since Otto Warburg considered ‘respiration damage’ to be a pivotal feature of cancer cells. Numerous somatic mitochondrial DNA (mtDNA) mutations have been found in various types of neoplasms, including breast cancer. Studies have shown that TNBC cells have profound metabolic alterations characterized by decreased mitochondrial respiration and increased glycolysis. Due to their impaired mitochondrial function, TNBC cells are highly sensitive to glycolytic inhibition, suggesting that such metabolic intervention may be an effective therapeutic strategy for this subtype of breast cancer cells [42]. Oxidative phosphorylation (OXPHOS) is an active metabolic pathway in many cancers. RNA from pretreatment biopsies from patients with triple-negative breast cancer (TNBC) who received neoadjuvant chemotherapy demonstrated that the top canonical pathway associated with worse outcome was higher expression of OXPHOS signature [43]. Henceforth, our EMT model TNBC transcriptomic analysis by IPA have selected the above mentioned, three very important canonical pathways involved in TNBC cell signaling.

Our transcriptomic analysis of the CRISPR-CAS9 genome edited Vimentin-RFP knock in TNBC cell line deciphered several differentially regulated genes and pathways that are involved in the EMT of these highly malignant breast cancer cells which could be of use for both diagnostic, prognostic, and therapeutic targets for future drug design and development.

## ACKNOWLEDGEMENT

The authors are grateful to Dr. Abedin of PRIMBIO Corporation for RNA Sequencing and data analysis.

This research was supported by NIH Grant# T34-GM100831, NSF-NOYCE Graduate student training award and a US Department of Education Graduate student training award to Elizabeth City State University Campus of The University of North Carolina.

